# Lichen fungi do not depend on the alga for ATP production

**DOI:** 10.1101/2021.03.17.435722

**Authors:** Gulnara Tagirdzhanova, John P. McCutcheon, Toby Spribille

## Abstract

Lichen fungi live in a symbiotic association with unicellular phototrophs and have no known aposymbiotic stage. A recent study postulated that some of them have lost mitochondrial oxidative phosphorylation and rely on their algal partners for ATP. This claim originated from an apparent lack of *ATP9*, a gene encoding one subunit of ATP synthase, from a few mitochondrial genomes. Here we show that while these fungi indeed have lost the mitochondrial *ATP9*, each retain a nuclear copy of this gene. Our analysis reaffirms that lichen fungi produce their own ATP.

## Introduction

In obligate symbioses, co-evolution of the partners often drives gene loss that results in complementarity of the symbionts’ metabolic capacities (e.g., Bublitz et al., 2019). Lichens are a diverse group of fungal-algal symbioses composed of at least one phototrophic partner (a green alga or a cyanobacterium) and at least one fungus. The fungus is currently assumed to be obligatorily associated with the phototroph. However, despite early suggestions for complementarity between fungal and phototroph gene products (Ahmadjian, 1993), evidence for this has been lacking. In 2018, Pogoda and colleagues were the first to report ostensible gene loss and complementarity in the lichen symbiosis. Based on analysis of mitochondrial genomes of several lichen-forming lecanoromycete fungi, Pogoda et al. (2018) reported that *ATP9*, a gene encoding F_1_F_0_ ATP synthase subunit C, one of the key proteins involved in oxidative phosphorylation, was missing from several fungal mitochondrial genomes (see also Funk et al., 2018; Stewart et al., 2018; Pogoda et al., 2019). For some of these species, the authors were able to find a copy of this gene in the nuclear genome (a gene transfer phenomenon known from a variety of ascomycetes, see Déquard-Chablat et al., 2011). For four lichen symbioses—*Alectoria fallacina, Gomphillus americanus, Heterodermia speciosa*, and *Imshaugia aleurites*—they did not detect any copy of the fungal *ATP9* gene. The authors concluded that in these symbioses, the fungus may rely on the alga for ATP production. This result has been since cited as evidence of obligate dependence of lichen fungi on their algal partners (e.g., Funk et al., 2018; Puri et al., 2021).

Several lines of evidence make this scenario improbable:

1. The complete loss of oxidative phosphorylation would inevitably be reflected in massive change in the mitochondrial genome (e.g., Heinz et al., 2012). The fact that all but one of the analyzed mitochondrial loci were found in all the genomes suggests that the function of mitochondria remains intact.
2. Fungal sexual reproduction via ascospores is intact in all four species; *Gomphillus americanus* reproduces only sexually. No vertical transmission is associated with this route. The ascospore has to be autonomous in order to germinate and find a compatible alga.
3. Close relatives of some of these species have been isolated in axenic cultures (e.g., *Heterodermia pseudospeciosa* and *Alectoria ochroleuca*; Crittenden et al., 1995; Yoshimura et al., 2002). They, therefore, are autonomous in ATP production.
4. All known instances of symbionts importing host ATP are from intracellular endosymbioses (e.g., Haferkamp et al., 2006). In lichens, the transfer would require sophisticated new mechanisms, given that ATP would need to move through the cell walls and membranes of both of the partners involved in the exchange.

We therefore hypothesized that the *ATP9* gene was present in the genomes but overlooked during the analysis. By replicating Pogoda et al. (2018) analysis on the species of interest, and then applying a series of stress tests, we were able to detect a putative homologue *ATP9* in all four fungi.

## Methods

### Sample preparation and sequencing

We generated metagenomic libraries for four lichen specimens: *Alectoria fallacina*, *Gomphillus americanus, Heterodermia speciosa*, and *Imshaugia aleurites* (Table S1). The samples were frozen at –80°C and ground in a TissueLyser II (Qiagen). We extracted DNA using QIAamp DNA Investigator Kit (Qiagen) for *Gomphillus* and DNeasy Plant Mini Kit (Qiagen) for the rest of the samples. The metagenomic libraries were prepared using Nextera Flex DNA kit (Illumina) and sequenced at the BC Cancer Genome Sciences Centre on an Illumina HiSeq X using 150 bp paired-end reads.

### Metagenomic assembly and genome annotation

The metagenomic data were filtered and assembled with the metaWRAP pipeline v1.2 (Uritskiy et al., 2018). We used the READ_QC module to remove any human contamination, and then assembled the remaining reads into metagenomes using metaSPAdes default settings (v3.13, Nurk et al., 2017). We binned individual assemblies using CONCOCT within metaWRAP (Alneberg et al., 2014). To identify the lecanoromycete genome assemblies among the bins, we analyzed each bin with BUSCO (v4.0.1, Seppey et al., 2019).

Some lecanoromycete genomes are heterogeneous in their GC content, which can result in these genomes being split between multiple bins (Tagirdzhanova et al., 2021). To obtain full genomes of the lecanoromycetes, we merged multiple bins as described in Tagirdzhanova et al. (2021). Briefly, we made GC-content vs coverage scatter plots for each metagenome and located the bin identified as an ascomycete genome by BUSCO. In all metagenomes except that of *Gomphillus*, these bins were part of a linear-shaped cloud (Fig. 1). In each metagenome individually, we merged bins forming this cloud into one MAG and confirmed with BUSCO that the merging improved completeness of the genome while maintaining low contamination.

**Fig. 1.**
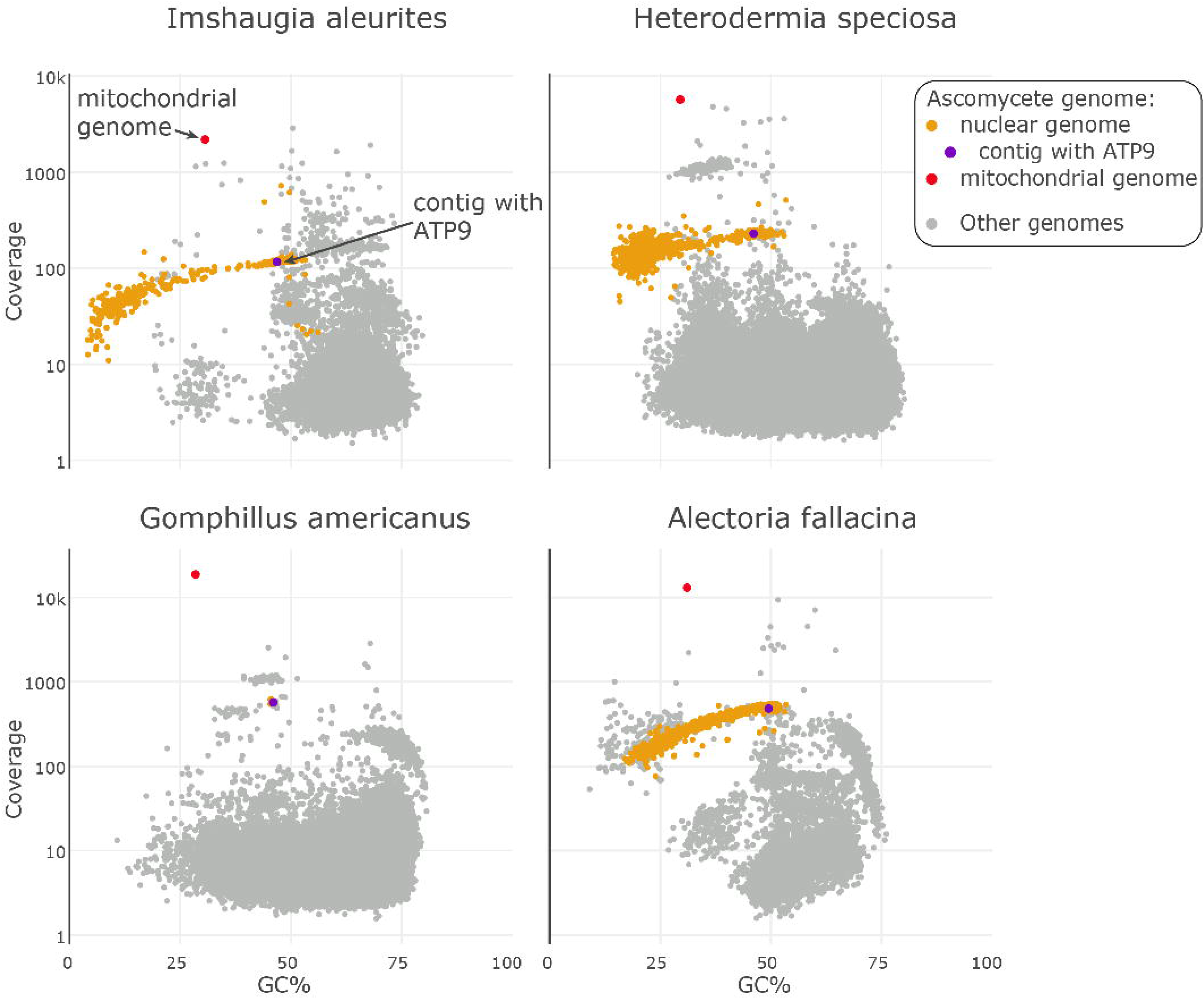
GC-coverage plots for the four metagenomes produced in this study. Dots representing contigs are positioned according to their GC content and coverage. Orange dots are contigs assigned to the lecanoromycete MAGs, purple dots are the contigs that contain the putative ATP9 homolog. Red dots are putative mitochondrial genomes.

We annotated the MAGs and six lecanoromycete genomes from GenBank (Table S2) using the Funannotate pipeline (v1.5.3, github.com/nextgenusfs/funannotate). We removed repetitive contigs from the assemblies, then sorted the assemblies and masked the repeats. Ab initio gene prediction was run using GeneMark-ES (v4.38, self-trained, Lomsadze et al., 2014), AUGUSTUS (v3.3.2, Stanke et al., 2004), SNAP (v 2006-07-28, Korf, 2004), and GlimmerHMM (v3.0.4, Majoros et al., 2006), trained using BUSCO2 gene models. We used EVidenceModeler (v1.1.1, Haas et al., 2008) to create consensus gene models, and removed models shorter than 50 amino acids or identified as transposons. The details on how we used funannotate are at github (https://github.com/metalichen/).

### Replicating Pogoda et al. (2018)

We searched the metagenomic assemblies using command line tBLASTn with default settings (v2.4.0, Camacho et al., 2009) and *ATP9, ATP8*, and *ATP6* genes from the mitochondrial genome of *Peltigera dolichorrhiza* as a query (NCBI Protein YP_009316289, YP_009316290, YP_009316291).

### Protein Family Dataset Assembly

We searched a recently published annotated genome of *Alectoria sarmentosa* (ENA GCA_904859925) for the genes assigned to pfam accession PF00137 and Interprosan accession IPR000454. We used the identified sequence as a query to locate putative lecanoromycete *ATP9* in the protein coding predictions produced by the genome annotation. We aligned all de-novo produced *ATP9* sequences against published sequences (Table S2) and manually curated the annotation. In case of three gene models *(Alectoria fallacina, Evernia prunastri*, and *Ramalina intermedia*; Table S3), we moved the intron boundaries to better match published *ATP9* sequences.

We extracted the putative protein sequences, and combined them with publicly available sequences of F_1_F_0_ ATP synthase subunit c from a variety of fungi and bacteria (Table S2). The sampling of the nuclear *ATP9* was done following Déquard-Chablat et al. (2011). As an outgroup we used N-ATPase following Koumandou & Kossida (2014). We aligned the sequences using MAFFT v7.271 (Katoh et al., 2002) with the flags --genafpair --maxiterate 10000 and excluded positions with more than 90% of data missing using trimal v1.2rev59 (Capella-Gutiérrez et al., 2009). The phylogeny was reconstructed with IQTree v1.6.12 (Nguyen et al., 2015) using LG+F+G4 substitution model and 50000 rapid bootstrap replicates.

### dN/dS analysis

To calculate the ratio of non-synonymous to synonymous substitutions (dN/dS) we followed Aylward (2018). We aligned the protein sequences of nuclear *ATP9* from lecanoromycetes using MAFFT as described above. We used this alignment together with the nucleotide sequences to create codon-based alignment with PAL2NAL (Suyama et al., 2006). To calculate the dN/dS ratios, we used codeml function in the PAML package (Yang, 2007).

## Results

### Pogoda et al. (2018) results replicated

We were able to replicate results of Pogoda et al. (2018) on our data. When using mitochondrial loci from the lecanoromycete of the *Peltigera dolichorrhiza* lichen as a tblastn query, we located *ATP6* and *ATP8*, but not *ATP9*. In all four metagenomes, *ATP6* and *ATP8* resided together in a single high-coverage contig (Fig. 1). The search for the gene in question, *ATP9*, failed to produce a blast hit above the threshold used by Pogoda et al. 2018 (bit score > 100).

### Putative ATP9 in the nuclear genomes

To test the hypothesis that the four species which Pogoda et al. reported as lacking *ATP9* in fact retain the gene, we began with the recently published lecanoromycete genome of *Alectoria sarmentosa* (Tagirdzhanova et al., 2021), a close relative of *A. fallacina*, one of the four fungi reportedly lacking *ATP9*. We identified one putative *ATP9* homologue, ASARMPREDX12_000654, in the *A. sarmentosa* lecanoromycete nuclear genome. This was the only gene from this genome assigned to Interproscan accession IPR000454 (ATP synthase, F_0_ complex, subunit C), and one of four assigned to pfam accession PF00137 (ATP synthase subunit C). When blasted against the NCBI Protein, it aligned with other fungal *ATP9* (Table S4).

Next, we generated metagenomes from newly acquired samples of all four lichen symbioses in which Pogoda et al. (2018) claimed fungal *ATP9* had been lost, and from them assembled and binned near-complete lecanoromycete genomes (metagenome-assembled genomes, MAGs).

Using ASARMPREDX12_000654 as a blast query, we found putative *ATP9* homologs in all MAGs. Each of these *ATP9* homologs showed up in the blast search we ran replicating Pogoda et al. (2018; see the previous section). However, their bit scores ranged from 35 to 48 and therefore were below the threshold set by Pogoda et al. (2018). We then checked the original metagenomic assemblies used in Pogoda et al. (2018) for the presence of these genes. Using the putative *ATP9* genes as a blast query we found similar genomic regions in all four genomes. For *Alectoria fallacina* and *Gomphillus americanus* the putative *ATP9* genes were identical in our assemblies and the assemblies from Pogoda et al. (2018); in *Heterodermia speciosa* and *Imshaugia aleurites* the sequences were > 98% identical with bit score > 1000.

Analysis of coverage suggests that the putative *ATP9* copy was located in the nuclear genome. In all four cases, contig coverage was similar to other contigs assigned to their respective MAGs and much less than that of the mitochondrial contig (Fig. 1). Of the six additional lecanoromycete genomes we surveyed, five contained putative nuclear *ATP9* (Table S2). In one of them, *Ramalina intermedia*, the nuclear *ATP9* homolog existed alongside the already reported *mtATP9* (NCBI Protein YP_009687549.1). Only in *Cladonia macilenta* were we unable to detect nuclear *ATP9*, but a fungal *mtATP9* was present.

### Two nuclear ATP9 homologs present in different Lecanoromycetes

We constructed a phylogeny of lecanoromycete *ATP9* genes identified in this study together with other fungal and bacterial *ATP9* genes. In the phylogeny, the putative lecanoromycete nuclear *ATP9* genes grouped together with known nuclear *ATP9* from other fungi (Fig. 2). The nuclear *ATP9* were split between two clades corresponding to *ATP9-5* and *ATP9-7* homologs described in Déquard-Chablat et al. (2011). All but one lecanoromycete nuclear *ATP9* were assigned to the *ATP9-5* clade; these fungi were from the Lecanoromycetes subclass Lecanoromycetidae. The only member of subclass Ostropomycetidae, *Gomphillus americanus*, grouped with *ATP9-7*.

**Fig. 2.**
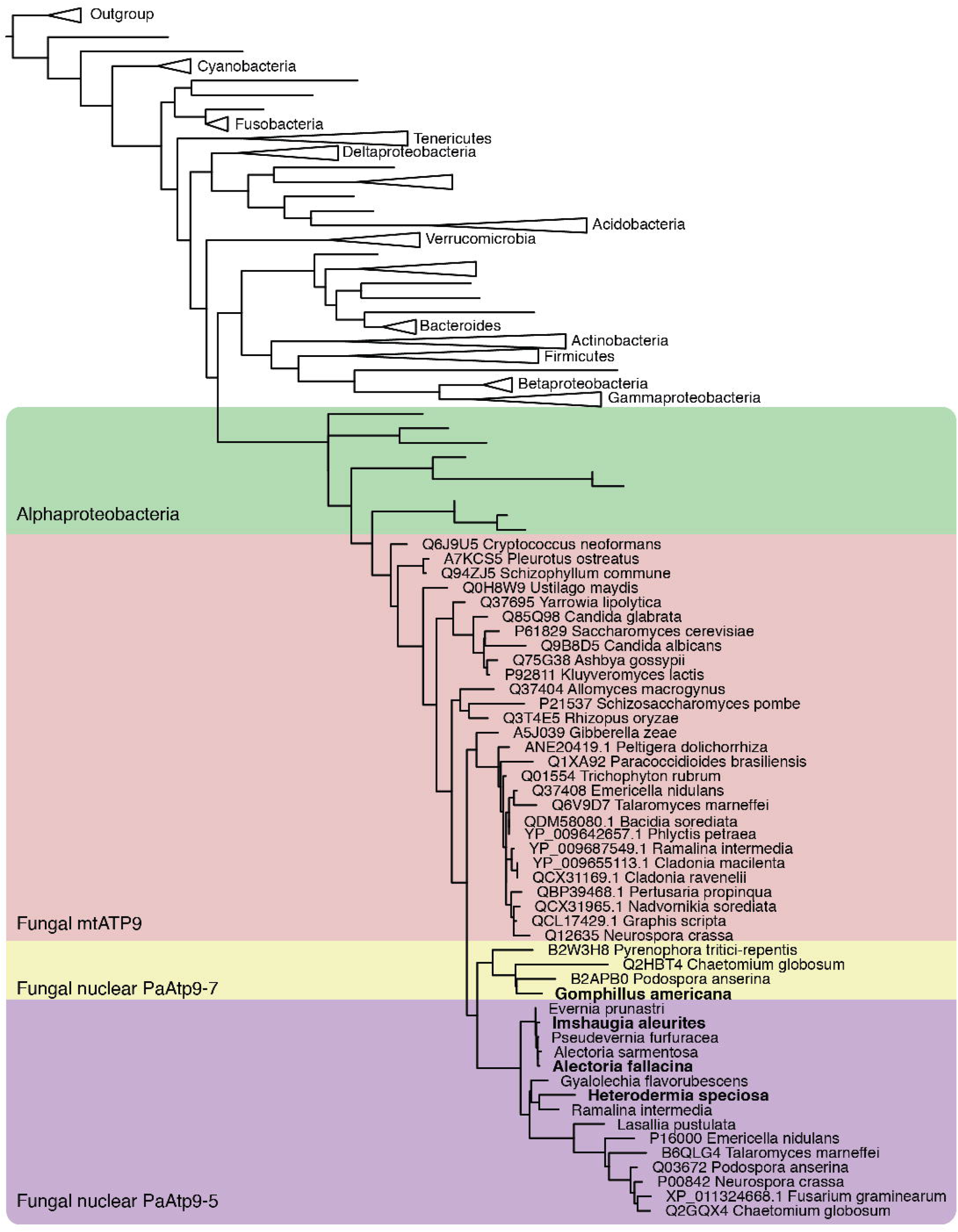
Phylogenetic tree of F_1_F_0_ ATP synthase subunit C across fungi and bacteria. ATP9 from the studied genomes are in bold.

The ascomycete nuclear *ATP9* clade was nested within the fungal *mtATP9*; its sister clade was formed by *mtATP9* from Pezizomycotina. This differs from the tree produced by Déquard-Chablat et al. (2011), as in their analysis the split between *ATP9-5* and *ATP9-7* is deeper and the two clades branch off in different places of the fungal *mtATP9* clade.

### Nuclear *ATP9* contain introns and are under purifying selection

All four putative *ATP9* contained at least one intron. In the three members of Lecanoromycetes subclass Lecanoromycetidae—*Alectoria fallacina*, *Heterodermia speciosa*, and *Imshaugia aleurites—ATP9* contained one intron, always in the same position (Table S3). The *Gomphillus americanus ATP9*, by contrast, contained two introns. The introns had either canonical GT-AG or one of the more common fungal non-canonical splicing sites (Table S3; Frey & Pucker, 2020). The dN/dS ratios between the nuclear *ATP9* from Lecanoromycetes ranged from 0.007 to 0.249 indicating that the gene is under purifying selection and is not a non-functional mitochondrial to nuclear genome transfer (see Richly & Leister, 2004).

## Discussion

Pogoda et al. (2018) hypothesized that some lichen fungi rely on other members of the symbiosis for ATP production based on the apparent lack of the *ATP9* gene in four Lecanoromycetes. We were able to find a putative *ATP9* homolog in all four genomes, both in new data produced for this study and in metagenomic data from the original publication. Our reanalysis reaffirms that, as expected, the fungi postulated to lack *ATP9* retain a nuclear copy of the gene, as in many other fungi. The fact that the putative *ATP9* were under purifying selection suggests that these genes are functional.

Our analysis suggests the nuclear *ATP9* originates in a transfer from the mitochondria to the nucleus, supporting the conclusion made by Déquard-Chablat et al. (2011). We included bacterial *ATP9* counterparts in the phylogeny to test for alternative hypothesis that the nuclear homologs are acquired not from mitochondria but from bacteria via horizontal gene transfer. This hypothesis was not supported: nuclear *ATP9* clade was nested within the *mtATP9* clade, which in turn was nested within Alphaproteobacterial clade.

Both known nuclear *ATP9* homologs, *ATP9-5* and *ATP9-7*, were present in the Lecanoromycete genomes. Déquard-Chablat et al. (2011) believed these genes to come from two independent transfers. They were previously reported in different combinations from several other classes of Pezizomycotina: Eurotiomycetes, Sordariomycetes, and Dothideomycetes (Déquard-Chablat et al., 2011). Adding Lecanoromycetes to the list further supports the hypothesis that the acquisition of *ATP9-5* and *ATP9-7* happened early in the evolution of Pezizomycotina and was followed by gene loss in some lineages.

With the combined evidence from this study and from Pogoda et al. (2018) we can begin to chart the evolutionary history of the *ATP9* in Lecanoromycetes. Most notably in the context of this study, several groups of Lecanoromycetes have lost *mtATP9* and retained only a nuclear copy.

We agree with Pogoda et al. (2018) in their assessment that the loss of *mtATP9* happened at least three times independently in the evolution of lecanoromycetes (see Fig. 1A in their study).

Gene loss affected nuclear *ATP9* homologs as well. None of the ten surveyed species retained both *ATP9-5* and *ATP9-7*: *Cladonia macilenta* had neither (while retaining *mtATP9*), the other species had either one or the other. Members of Lecanoromycetidae, other than *Cladonia*, retained *ATP9-5*, while the only member of Ostropomycetidae retained *ATP9-7*. Further research will map the nuclear *ATP9* across the lecanoromycete fungi and check how the new data points alter our understanding of the evolutionary history of this gene.

Our reanalysis of the Pogoda et al. (2018) paper underlines that the apparent lack of any one gene does not automatically translate into the loss of biological function, especially when the rest of the pathway is maintained. While *ATP9* indeed appears missing from mitochondrial genomes of some Lecanoromycetes, this result by itself was not sufficient to back the claim of lichen fungi having lost oxidative phosphorylation.

## Acknowledgments

We thank the authors of the original publication for access to their data. We thank Jason Hollinger and Dylan Stover for providing specimens from which we sequenced metagenomes, and David Díaz-Escandón and Sophie Dang for helping with lab work. GT and TS were supported by an NSERC Discovery Grant and a Canada Research Chair in Symbiosis.

## Author Contributions

G.T., T.S., and J.M. designed the study and wrote the manuscript. G.T. gathered and analyzed the data and produced figures.

## Data Accessibility Statement

Raw metagenomic data, metagenomic assemblies, and MAGs: European Nucleotide Archive (PRJEB42325). Custom scripts: https://github.com/metalichen/Lichen-fungi-do-not-depend-on-the-alga-for-ATP-production.

## Notes

### Competing Interest Statement

The authors have declared no competing interest.

